# ELAplus: Fast and accurate analysis platform for energy landscape analysis facilitated by fine-tuning optimization algorithm

**DOI:** 10.64898/2026.08.26.747178

**Authors:** Sotaro Takano, Hiroaki Fujita, Fumihiko Ayabe, Zeus Sato, Hiroshi Masuya, Hirokazu Toju, Kenta Suzuki

## Abstract

1. Large-scale community-composition datasets, especially from microbiome studies, increasingly provide opportunities to identify major community compositional types (e.g., enterotypes in human microbiomes) and potential transitions depending on environmental factors. Energy landscape analysis based on maximum entropy models has emerged as a promising framework for characterizing such multi-stability in ecological communities. However, its application to diverse, high-dimensional compositional datasets remains limited by computational inefficiency, insufficient evaluation of predictability, and lack of systematic assessment of uncertainty.
2. Here, we present a computationally tractable inference framework for energy landscape analysis of multispecies communities, implemented in the R package ELAplus. We introduce a framework combining cross-validation-based selection of optimization settings, enabling accurate and computationally efficient model fitting across a wide range of simulated community datasets. In addition, we incorporate a bootstrap-based approach to quantify the reliability of inferred stable states, providing a systematic measure of uncertainty in landscape structures.
3. Simulation analyses demonstrate improved predictive performance and robustness compared to existing implementations. Applications to simulated occurrence datasets further illustrate how the framework can reveal stable states, basins of attraction, and potential tipping points under varying environmental conditions. The package also provides visualization tools, including disconnectivity graphs and energy surface plots, to facilitate intuitive interpretation of complex ecological landscapes.
4. Our framework enables robust and computationally efficient inference of ecological stability from compositional and environmental data, expanding the applicability of energy landscape approaches in diverse natural communities.

## 1. Introduction

Across diverse ecosystems, ecological communities can undergo abrupt regime shifts between alternative states, with consequences for ecosystem functioning, climate regulation, and biogeochemical cycling (Scheffer *et al*. 2001; Flores *et al*. 2024). Analogous patterns of distinct community composition have recently attracted attention in microbiome research, notably human gut enterotypes and disease-associated dysbiosis (Arumugam *et al*. 2011; Costea *et al*. 2018). However, enterotypes are empirically identified compositional clusters that possibly arise from environmental variation or community assembly processes (Gonze *et al*. 2017); whether they represent alternative stable states and how transitions occur therefore remain unclear. Energy landscape analysis (ELA) is a data-driven approach and addresses this gap by inferring potential stable states and transition structures from multivariate observational datasets. The concept of ELA have been originally developed to model neural dynamics in the brain (Watanabe *et al*. 2014; Masuda *et al*. 2025) and recently extended to ecological community data, enabling the inference of alternative stable states and their transition structures across environmental gradients from species occurrence patterns (Suzuki *et al*. 2021).

Energy landscape of ecological communities can be generally conceptualized by a graphical network (`ELA()` panel in Figure 1), where nodes denote community compositions and edges represent transitions between adjacent compositions (i.e., presence/absence status of the one community member changes). In our framework, ELA formulates the probability of each community composition by extended pairwise maximum entropy model (`runSA()` panel in Figure 1, EPMEM hereafter) that incorporates pairwise taxon relationships and environmental effects. EPMEM identified least-biased distribution consistent with the observed occurrence datasets based on the maximum entropy principle in statistical physics (Jaynes 1982). This estimated probability, in turn, corresponds to the inverse proportion metric of energy in the principle of maximum entropy (i.e., the lower energy in a community composition is indicative of higher probability). Then, the implementation of estimated energy value at each node on the graph enables the representation of the probability of transitions between community compositions as a landscape (`ELA()` panel in Figure 1). In addition, EPMEM enables to reconstruct the changes in the landscapes across environmental gradients by integrating environmental parameters in the model, capturing environment-induced regime shifts (`gradELA()` panel in Figure 1).

**FIGURE 1.**
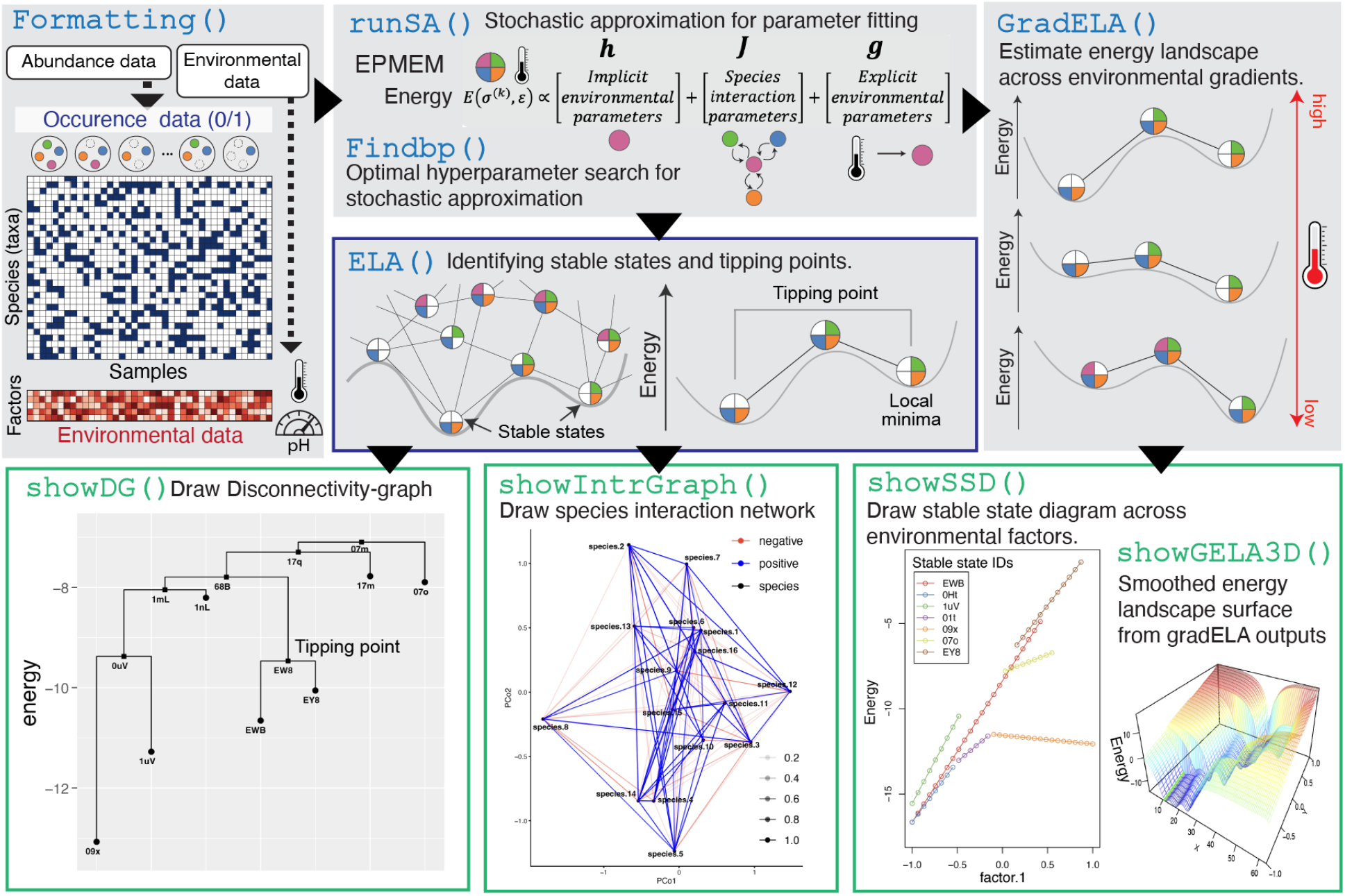
Overview and important functions of the ELAplus R package. The workflow starts from formatting of input datasets followed by parameter fitting, construction of the landscape (encircled by blue), the landscape construction across environmental parameters, and then visualization (encircled by green). The title of each panel corresponds to the R function name for each process.

The ELA framework has recently been applied to diverse ecological datasets to infer potential basins and forecast abrupt shifts in community composition (Fujita *et al*. 2023; Kadoya, Suzuki & Terui 2025; Toju *et al*. 2026).However, characteristic features of ecological datasets pose distinct challenges for EPMEM fitting. One such characteristic in the dataset is high-dimensionality. For instance, microbiome data usually retain tens to hundreds of taxa even after taxonomic aggregation, substantially increasing the number of model parameters to be estimated.. In such cases, conventional gradient descent approaches, as used in early applications to neural activity data (Watanabe *et al*. 2014), are computationally expensive, motivating the use of a more scalable alternative. Stochastic maximum likelihood (SML), approximating model expectations through Monte Carlo sampling, can addresses computational aspect and recently applied to the ecological datasets (Suzuki *et al*. 2021). Another characteristic is sparsity. Ecological occurrence matrices often contain many absent taxa, making pairwise relationships difficult to estimate reliably (Weiss *et al*. 2016). Consequently, EPMEMs possibly infer weakly supported pairwise associations and overfit the observed data. Because ecological association networks are often expected to contain only a subset of all possible pairwise links (Busiello *et al*. 2017; Weiss *et al*. 2022), L1 regularization can mitigate this problem by promoting a sparse interaction network (Cheng *et al*. 2014). The accuracy and convergence of SML-based fitting depend on the hyperparameters such as the learning rate and the strength of L1 regularization (Berglund & Raiko 2013; Leclerc & Madry 2020; Liu, Gao & Yin 2020; Fu *et al*. 2023). These considerations highlight the need for a systematic control of the optimization process by selecting dataset-specific optimization configurations.

Here, we developed ELAplus, a R package that provides a platform for analysis and visualization of energy landscape. There are notable features for ELAplus, (i) We developed cross-validation method to select dataset-specific optimization settings and stopping iterations (Table 1), together with L1 regularization to reduce overfitting in sparse occurrence data, thereby improving the accuracy and reliability of EPMEM fitting across diverse community datasets. (ii) To enhance the intuitive interpretation of multi-stability, we developed and implemented the built-in visualization functions for key properties in the landscape, such as stable states, basins, and tipping points (Table 2) through disconnectivity graphs and surface plots. (iii) To improve the robustness of inferred energy landscapes, we developed pruning method, which removes shallow basins by merging them into neighboring deeper basins according to their energy barriers.

**TABLE 1.**
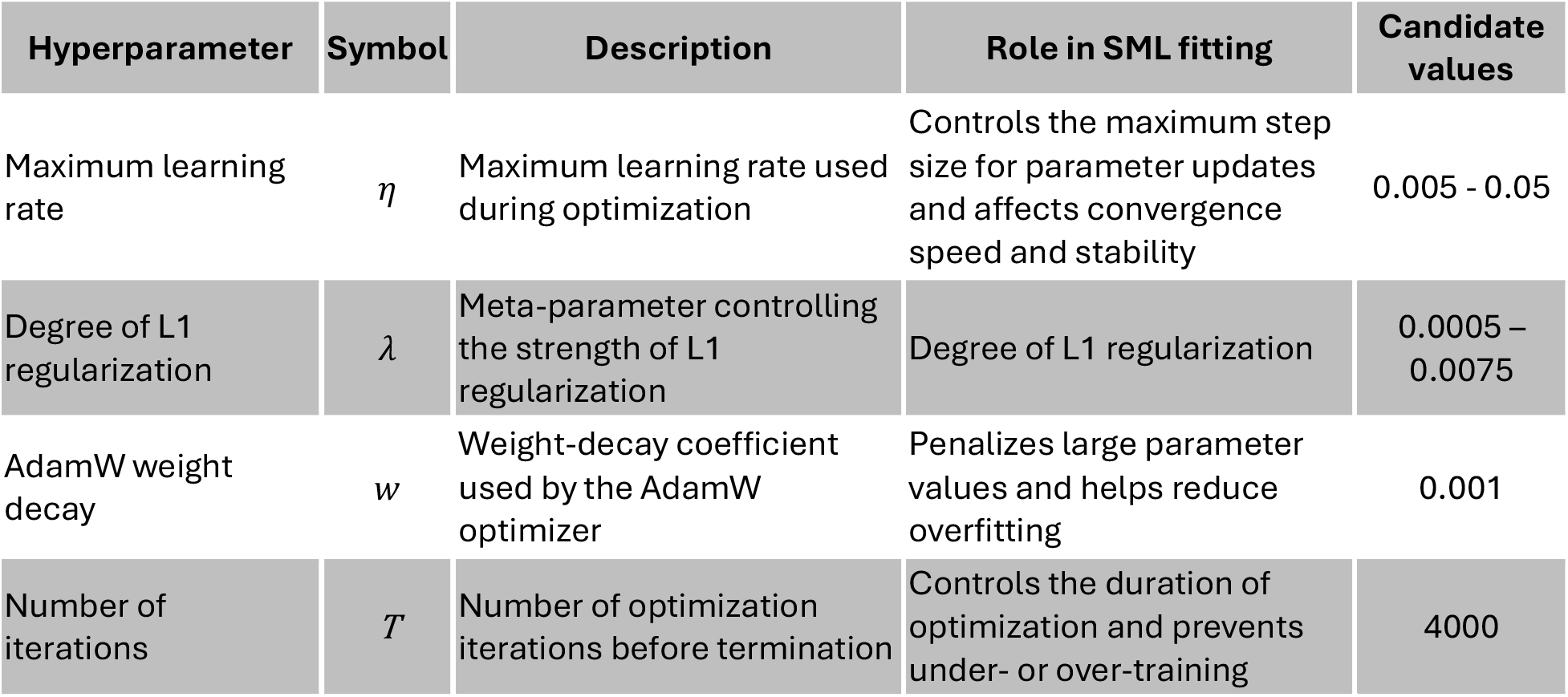
Optimization settings used in the model fitting procedure of EPMEM.

| Hyperparameter | Symbol | Description | Role in SML fitting | Candidate values |
| --- | --- | --- | --- | --- |
| Maximum learning rate | $\eta$ | Maximum learning rate used during optimization | Controls the maximum step size for parameter updates and affects convergence speed and stability | 0.005 - 0.05 |
| Degree of L1 regularization | $\lambda$ | Meta-parameter controlling the strength of L1 regularization | Degree of L1 regularization | 0.0005 – 0.0075 |
| AdamW weight decay | $w$ | Weight-decay coefficient used by the AdamW optimizer | Penalizes large parameter values and helps reduce overfitting | 0.001 |
| Number of iterations | $T$ | Number of optimization iterations before termination | Controls the duration of optimization and prevents under- or over-training | 4000 |

**TABLE 2.** Key properties in ELA.

| Term | Definition and meaning | Ecological interpretation |
| --- | --- | --- |
| Community state | A community composition represented by the presence or absence of individual taxa, typically encoded as a binary vector. | A possible taxonomic composition of the community. |
| Energy | A potential value assigned to each community state. | States with lower energy are interpreted as being relatively more probable to maintain or appear. |
| Stable state | A community state with local energy minima. It corresponds to the bottom of a basin (see ELA ( ) in Figure 1). | A representative community composition that is relatively persistent. This is not a typical dynamical equilibrium or attractor. |
| Basin | A set of community states that reach the same stable state through steepest-descent transitions. | A range of community compositions that tend to converge to the same representative stable state. |
| Tipping point | The lowest-energy boundary state between two stable states (see ELA ( ) in Figure 1). | A critical intermediate community composition associated with a transition from one stable state to another. |
| Disconnectivity graph | A hierarchical graph summarizing stable states and the minimum energy barriers separating them. | Provides an overview of the transitions between representative community compositions. |

We applied those approaches to the prediction of population dynamics in multispecies system and comprehensively estimated the applicability of the established method.

## 2. Outline of ELAplus package

In ELAplus, all analyses were implemented based on a same framework as previously formalized (Suzuki *et al*. 2021; Kadoya, Suzuki & Terui 2025) (for details, see Supporting information). The typical workflow consists of: (i) Formatting the datasets such as binarization of species (taxon) abundance matrix (ii) Fitting the species (taxon) occurrence data to EPMEM, (iii) constructing an energy landscape from the fitted model, (iv) Identifying stable states and transition structures, and (v) Visualizing the resulting landscape and interactions (Figure 1).

### 2.1 Formatting datasets

ELAplus accepts a sample-by-taxon abundance table and optional environmental metadata. The `Formatting()`function converts taxon abundances to relative abundances and then to binary occurrence data using a user-defined threshold. Taxa with very low or high occurrence frequencies can be filtered because they provide insufficient variation for reliable parameter estimation. When environmental variables, such as temperature or pH, are provided, they are matched to the corresponding community samples and formatted for subsequent EPMEM fitting (for details, see Supporting Information).

### 2.2 Parameter fitting to EPMEM

Formatted datases are fitted to the EPMEM by `runSA()` function. The EPMEM icludes three parameters (for details, see Supporting information): ***J***, the strengths of pairwise relationship between taxa; ***g***, the taxa-specific effect of environmental factors such as temperature and pH; ***h***, the intrinsic tendency of each taxon to occur that cannot be explained by pairwise relationships (***J***) or explicit environmental factors (***g***). We estimated the model expectation in the likelihood gradient using a persistent Gibbs sampling approach, where the sampled states were updated once during each parameter update step, usually known as stochastic maximum likelihood (SML) (for details, see Supporting Information). The `runSA()` function runs SML algorithms with the user-defined optimization settings. To avoid the unstable prediciton, we implemented `Findbp()` function in ELAplus, which provides a dedicated grid-search routine to choose hyperparameters and stopping criterion empirically (the impact of this process is explained in 3. Results section).

### 2.3 Construction of the energy landscape

After EPMEM fitting, `ELA()`function constructs energy landscape over community compositions and identified local energy minima (stable states), their basins, and tipping points (`ELA()` panel in Figure 1 and Table 2).

The inferred landscape is estimated from a finite set of observed datasets, which possibly generates shallow (statistically unreliable) basins due to sampling variability. `ELPruning()` function merges stable states separated by low energy barriers and removes potentially spurious stable states, thereby retaining the major multi-stable structure of the landscape, which results in a more robust and interpretable energy landscape (for details, see **3.5**).

The landscape of a community is not fixed, but changes continuously along an environmental gradient (e.g., temperature, nutrient availability, disturbance intensity). `GradELA()` is designed to capture this dependence by estimating energy landscapes repeatedly across a controlled range of an environmental factor and summarizing how stable states and transition barriers change.

### 2.4 Visualization

ELAplus provides complementary visualizations of the fitted model and landscape. `showDG()` function helps reveal the hierarchical organization of basins such as funnel-like structures by visualizing a disconnectivity graph. In the graph, terminal branches correspond to stable states (local minima), while branch merging heights represent the minimum energy barriers between states (tipping points). In ecological applications, transition barriers are directly related to transition probabilities and metastability, and disconnectivity graphs can be used to visualize alternative stable community states and the difficulty of transitions between them.

The fitted parameter ***J*** in the EPMEM enabled inference of the relationship strengths between taxa, and `showIntrGraph()` visualizes pairwise relationships among taxa and helps identify hub taxa within the interaction network.

When environmental data are included in the analysis, `showSSD()`and `showGELA3D()`can visualize the results of `gradELA()`, which illustrates regime shifts by showing how the number, identity, and energy of stable states vary along an environmental gradient. The two functions are conceptually similar, except that `showSSD()` produces a two-dimensional visualization, whereas `showGELA3D()` provides a three-dimensional representation (Figure 1).

## Results

### 3.1 Cross-validation approach for optimal parameter fitting

Fitting an EPMEM to observed data using SML is a major bottleneck in ELA in terms of both computation time and landscape accuracy. Although the optimization problem has a unique optimum when the exact gradient is available, the likelihood gradient must be approximated using SML for high-dimensional datasets. Fitting performance therefore depends on the optimization settings. Moreover, with finite and noisy observational data, excessive optimization may capture spurious structures in the training data (Berglund & Raiko 2013), reducing the stability and generalizability of the inferred landscapes. To address these issues, we implemented `Findbp()`function, which systematically searches for optimization settings that balance computational efficiency and fitting accuracy. `Findbp()`determines the total iterations by cross-validation based method (CV), in which the data were divided into training and validation sets, and training was stopped at the iteration where the validation loss (i.e., negative conditional log-likelihood of the occurrence matrix estimated by the trained EPMEM to the test data, for details see Supporting Information) reached its minimum (Figure 2C, CV). For comparison, we also evaluated a conventional convergence criterion (Figure 2B, CONV), where the optimization was terminated when the update statistic no longer decreased over a predefined number of iterations (Figure 2B, CONV; see also Eqs. (12) - (14) in Supporting Information). In this example, CONV continued optimization beyond the minimum validation loss (Figure 2C) because its stopping criterion depended solely on the parameter-update statistic. In contrast, CV reduced overfitting and improved parameter estimation. To evaluate parameter-estimation accuracy, we randomly generated EPMEM parameter sets (***h, J***, and ***g***) and used Gibbs sampling to simulate occurrence datasets with 48 taxa and 512 samples. The parameter values used to generate each dataset were treated as the *“*true*”* parameters and compared with their estimates using Spearman correlations. Across 512 randomly generated datasets, CV produced higher correlations between the true and estimated parameters than CONV (Figure 2D). Thus, CV reduced overfitting and provided a more reliable stopping criterion than numerical convergence alone.

**FIGURE 2.**
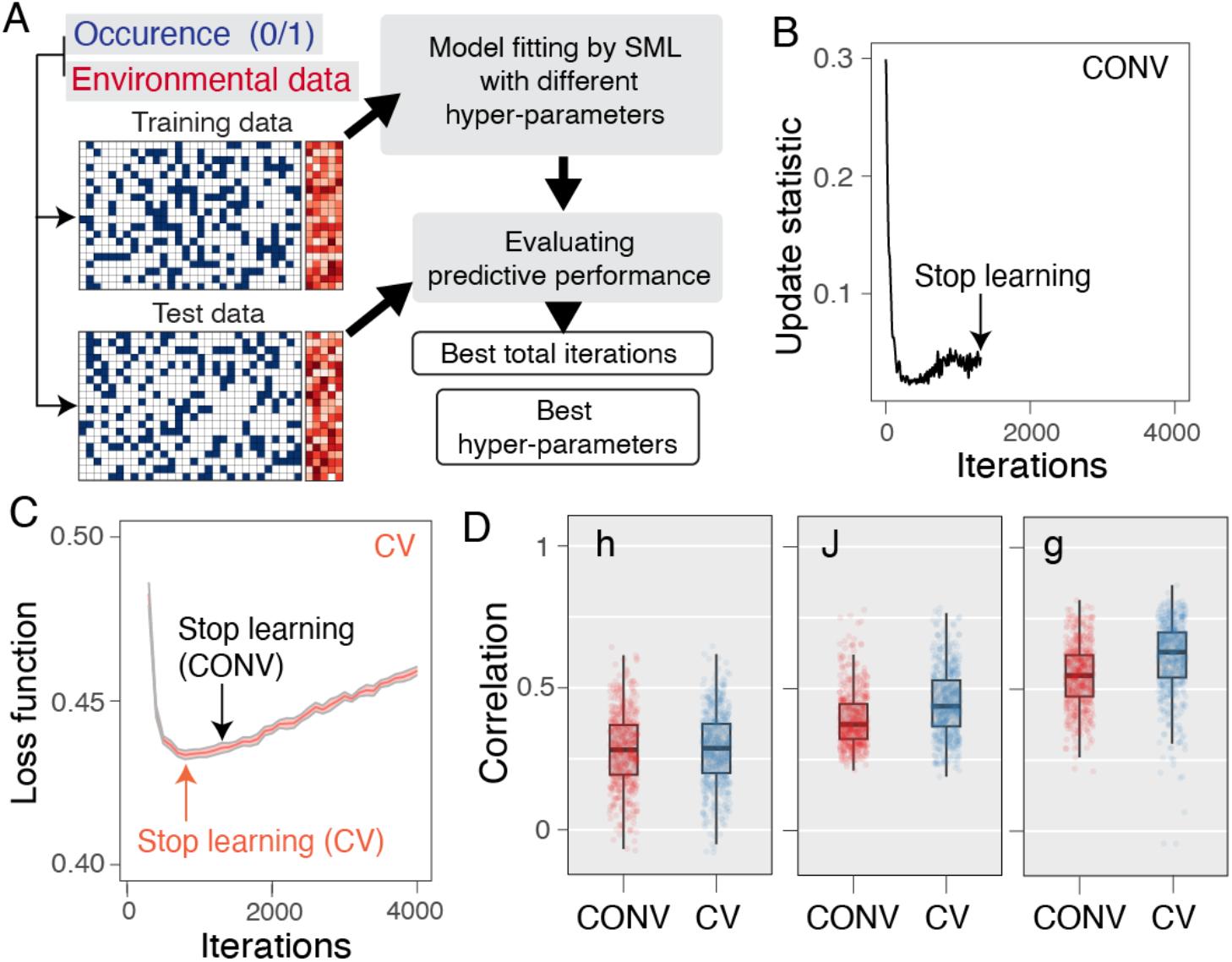
Cross-validation-based selection of optimization settings improves SML prediction. (A) Overview of `Findbp`(). The function generates training–test splits, evaluates combinations of hyperparameters and iterations by cross-validation, and selects the combination minimizing the prediction error for held-out samples. (B) Example changes in parameter-update magnitude during fitting. The fitting is terminated based on the convergence criterion (CONV). (C) Corresponding cross-validation loss for held-out test data. Black and red arrows indicate stopping points selected by the convergence criterion (CONV) and cross-validation (CV), respectively. (D) Spearman correlations between true and estimated EPMEM parameters for 512 randomly generated parameter sets, compared between stopping criteria. Bold lines indicate medians, and different letters denote statistically significant differences. Taxon–taxon interaction connectance, the proportion of nonzero off-diagonal elements in ***J***, was set to 0.5. Results are shown for datasets containing 48 taxa.

### 3.2 L1 regularization in parameter estimation for sparse ecological networks

Ecological occurrence matrices are typically sparse, and consequently only a subset of pairwise relationships is expected to contribute substantially to community assembly. To account for this property, ELAplus implements sparse parameter estimation by incorporating L1 regularization with cumulative weight penalties (for details, see Supporting Information) (Tsuruoka, Tsujii & Ananiadou 2009). For datasets generated from low-connectance interaction networks, L1 regularization consistently improved parameter estimation, particularly for the pairwise relationship matrices ***J*** (Figure 3A, Connectance *=* 0.2). In contrast, for datasets generated from more densely connected interaction networks (connectance = 0.5), L1 regularization alone resulted in lower predictive performance (Figure 3B), indicating that the effectiveness of sparse regularization depends on the underlying network connectance. Notably, this dependence was largely eliminated when L1 regularization was combined with the adaptive optimization algorithm described in Section **3.3**.

**FIGURE 3.**
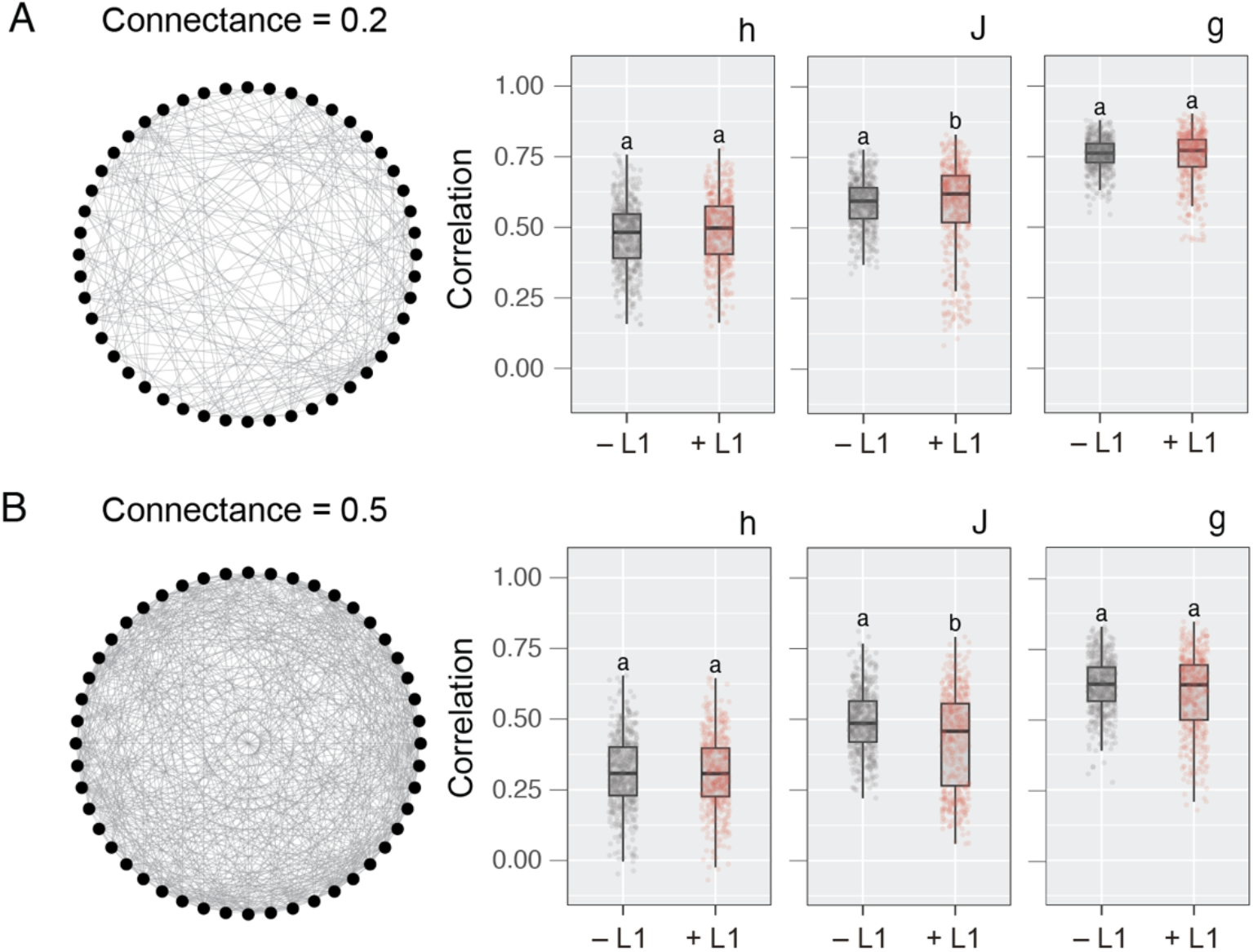
L1 regularization improves parameter estimation for occurrence datasets generated from interaction networks with different connectance. Spearman*’*s correlations between the true and estimated EPMEM parameter sets for 512 randomly generated datasets with low (A, connectance = 0.2) and high (B, connectance = 0.5) interaction-network connectance. Results obtained without and with L1 regularization are shown in gray and red, respectively. Bold lines indicate the median, and statistically significant differences are indicated by different letters. The results shown are based on datasets containing 48 taxa.

### 3.3 Adaptive optimization algorithm improves parameter-estimation performance

Next, we examined whether incorporating an adaptive optimization algorithm into SML-based EPMEM fitting improves parameter-estimation accuracy. We compared the simple momentum method (Qian 1999; Suzuki *et al*. 2021) with adaptive moment estimation with decoupled weight decay (AdamW) [8, 10]. Because AdamW generally converges more efficiently and with lower error than momentum (Kingma 2014; Zhang *et al*. 2020), we tested whether this advantage extends to EPMEMs using randomly generated training datasets (Figure 4). AdamW improved the predictive performance for all three parameters (Figure 4A), particularly for datasets with many taxa (Figure S1), supporting its use for large communities. However, the validation loss eventually increased during training (Figure 4B, black line), indicating overfitting despite the use of decoupled weight decay (Loshchilov & Hutter 2017).

**FIGURE 4.**
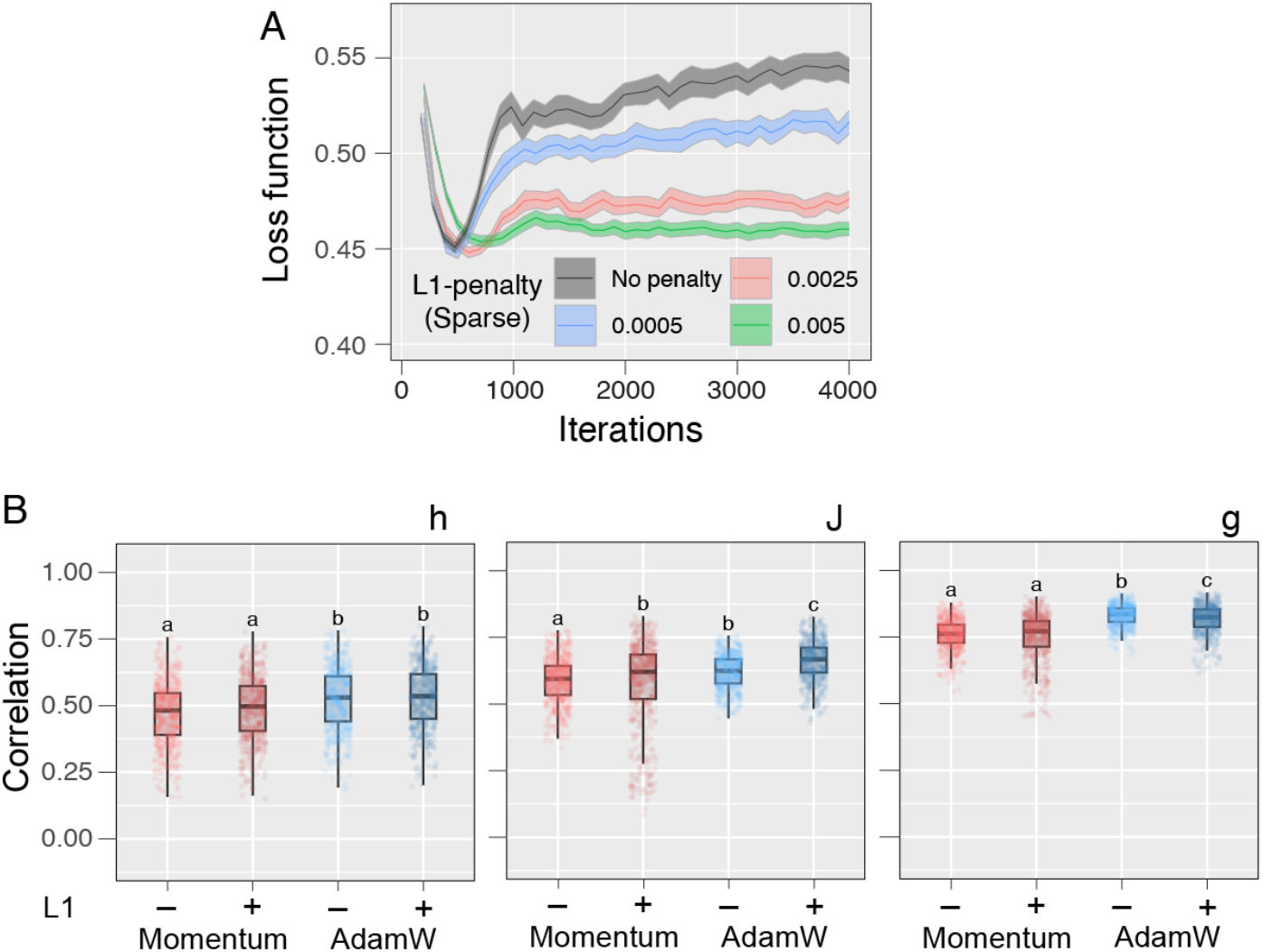
Combining AdamW and L1 regularization improve parameter estimation performance. (A) Example changes in the validation loss with AdamW and/or L1 regularization. Shaded regions indicate 95% confidence intervals across different training/test splits. (B) Spearman*’*s correlations between true and predicted parameter sets in EPMEM for 512 random parameter sets, showing the effects of AdamW and/or sparse modeling on predictive performance in SML. In panels B, bold lines indicate median values, and statistically significant groups are denoted by different letters. The connectance of taxa&taxa interactions (i.e., the proportion of nonzero off-diagonal elements in ***J***) in random datasets was set to 0.2. The results shown are based on datasets containing 48 taxa.

Overfitting can be mitigated by stopping training at the minimum validation loss (Figure 2) or by imposing sparsity on the interaction parameters through L1 regularization (Tibshirani 2011; Sheppert 2026). Combining L1 regularization with AdamW reduced overfitting (Figure 4B) and improved predictive performance, particularly for low-connectance interaction networks (Figure 4C; connectance = 0.2). For more densely connected networks (connectance = 0.5 or 0.8), this combination performed comparably to the unregularized model (Figure S2). Thus, L1 regularization and AdamW play complementary roles: together, they improve parameter estimation for sparse networks without compromising performance for densely connected networks. This broad applicability is advantageous because the connectance of ecological communities is generally unknown before analysis.

To determine whether improved EPMEM fitting translated into more accurate landscape inference, we compared ELA predictions with community states generated by competitive Lotka– Volterra simulations (Figure S3A and S3B). The use of AdamW and L1 regularization increased the recall of stable states and strengthened the correlation between the observed and predicted occurrence probabilities of community compositions (Figure S3C and S3E). However, these improvements were accompanied by reduced precision, indicating that the increased recovery of potential stable states came at the cost of additional false-positive predictions (Figure S3D). This trade-off motivated the subsequent bootstrap-based assessment of the reliability of predicted stable states (for details, see Supporting Information).

### 3.4 Bootstrap-based assessment of the reliability of inferred landscape

The outcome of SML can be usually influenced by variation in the observed community compositions used for parameter estimation. Assessing variation in predicted EPMEM parameters across resampled training datasets is therefore important for evaluating the robustness of the inferred landscape. Of particular interest is basin depth (Fig 5A, *ΔE*), defined here as the energy difference between a stable state and the lowest tipping point connecting it to another stable state. Ecologically, deeper basins correspond to community compositions that are more resilient to perturbations and therefore more likely to persist, whereas shallow basins indicate states that can transition more readily following compositional disturbances.

**FIGURE 5.**
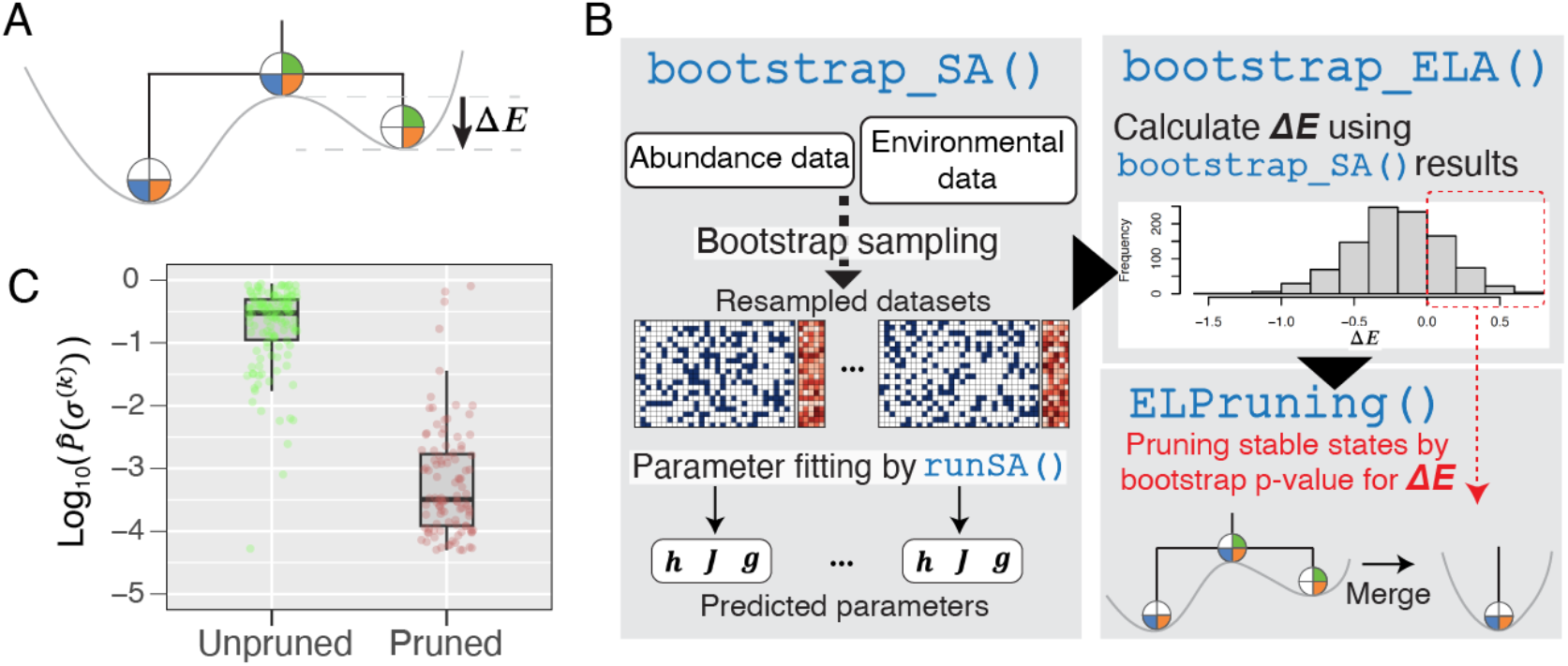
Bootstrap-based pruning improves stable-state prediction in the LV system. (A) Schematic illustration of the energy gap (*ΔE*) in the energy landscape. (B) Overview of the bootstrap-based pruning procedure for predicted stable states. The `bootstrap_SA()` function performs parameter fitting using bootstrap-resampled training datasets, and the resulting EPMEM models are used in `bootstrap_ELA()` to calculate the energies of stable states and tipping points. The resulting *ΔE* distributions are then used to estimate bootstrap-based p-values. In `ELPruning()`, stable states with high approximated p-values are pruned and merged into connected deeper stable states. (C) Comparison of the empirical appearance probabilities (i.e., observed probabilities in LV system) between unpruned (more stable) and pruned (less stable) stable states in ELA. Bold lines indicate median values, and statistically significant groups are denoted by different letters.

To assess how sampling variation in the training data affects inferred landscape structures, particularly the reliability of predicted stable states, ELAplus can generate bootstrap-resampled datasets and independently refits the EPMEM using `bootstrap_SA()`Figure 5B). Each bootstrap dataset is generated by resampling rows of the occurrence matrix with replacement while preserving the correspondence between species occurrence profiles and environmental variables, and contains the same number of samples as the original dataset. The resulting parameter estimates were used to calculate the distribution of energy gaps between the predicted stable states and their corresponding tipping points (Figure 5B, `bootstrap_ELA()`) as described previously (Suzuki *et al*. 2026). A statistically reliable stable state is expected to have Δ*E* < 0 consistently across bootstrap replicates. We therefore define the proportion of replicates with Δ*E* >= 0 as a *“*bootstrap-approximated p-value*”* with smaller values indicating greater statistical reliability. In the example case, more than 25% of bootstrap replicates had Δ*E* > 0, yielding p-value > 0.25). `ELPruning()`uses this measure to remove weakly supported stable states. For instance, when the threshold is set to `p-value = 0.25` in `ELPruning()`, the function iteratively selects the pair containing the shallowest basin among those for which either stable state has a p-value > 0.25 and merges the shallower state into the deeper one. This procedure is repeated until no pair meets the p-value criterion (for details, see Supporting Methods).

To evaluate whether the statistical reliability estimated by bootstrap analysis reflects the stability of inferred stable states in the empirical population dynamics, we compared the bootstrap results with the simulated dynamics by a Lotka–Volterra (LV) model (for details, see Supporting text). If the bootstrap-derived reliability reflects the stability of stable states in the underlying population dynamics, stable states with lower statistical support (i.e., large bootstrap-approximated p-value) should be visited less frequently and transition more readily to alternative states in the LV dynamics. We checked this possibility by examining the empirical appearance probability in the LV simulations 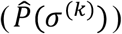, defined as the proportion of appearance that the LV trajectories occupied the corresponding stable state (σ^(*K*)^) during long-term simulations. The appearance probability of unpruned stable states are significantly higher than that of pruned stable states by `ELPruning()` (Figure 5C), indicating that the estimated statistical reliability corresponds to the stability in the simulated population dynamics.

## Conclusions

We developed ELAplus, based on extended pairwise maximum entropy models. By integrating cross-validation, AdamW optimization, and sparse regularization, the framework substantially improved parameter fitting accuracy and predictive performance, particularly in high-dimensional community datasets. Simulation analyses using competitive Lotka–Volterra systems demonstrated that these optimizations enhanced the recovery of stable states and the prediction of community stability landscapes. In addition, the bootstrap-based pruning framework provided a practical approach for evaluating the statistical reliability of inferred stable states and transition structures. Together with built-in visualization tools, ELAplus offers an accessible platform for investigating multistability, regime shifts, and transition dynamics in ecological systems. Our framework expands the applicability of energy landscape approaches to large-scale compositional and environmental datasets.

## Code availability

The ELAplus R package is available from CRAN (https://CRAN.R-project.org/package=ELAplus), and its source code is available in the GitHub repository (https://github.com/sotarotakano/rELA_CRAN).

## Author contributions

K.S. conceptualized the original framework. S.T., H.F., F.A., Z.S., and K.S. contributed to software development. S.T. contributed to formal analysis, visualization. and wrote the original draft. All authors, including S.T., H.F., F.A., Z.S., H.M., H.T., and K.S., contributed to data interpretation and the writing of the manuscript.

## Supporting information

Supporting text Fig. S1-S3

