## Supporting Information for "ELAplus: Fast and accurate analysis platform for energy landscape analysis facilitated by fine-tuning optimization algorithm"

### Supporting Text

#### Overview of ELA

In energy landscape analysis (ELA), the community composition, denoted by $\boldsymbol{\sigma}$, is represented by a binary vector of length $S$, the total number of species. There are $2^S$ unique community compositions in total, where each node in the network represents a presence–absence status of the community members (pie charts in `ELA()` section in Fig. 1). Each node connects to its all-possible adjacent nodes (i.e., presence-absence status (1/0) of only one specific member is reverted), resulting in a regular network with $S$ links per node. The binary composition vector $\boldsymbol{\sigma}$ always changes in a stepwise manner (Fig. 1).

Then, the likelihood of the occurrence of each node was computed by EPMEM based on species co-occurrence patterns and environmental parameters (Fig. 1). Here, we designate this likelihood as "energy" following the terminology of statistical physics. The EPMEM estimates the energy (occurrence probability) of the community composition $\boldsymbol{\sigma}^{(k)}$ in each environmental condition denoted as $\boldsymbol{\varepsilon} = (\varepsilon_1, \varepsilon_2, \varepsilon_3 \ldots, \varepsilon_M)$.

The EPMEM model consists of three parameters: $\boldsymbol{h}$, $\boldsymbol{g}$, and $\boldsymbol{J}$. $\boldsymbol{h}$ $(h_1, h_2, h_3, \ldots, h_S)$ represents the net effect of implicit environmental factors where the $i$th taxon is more likely to be present ($h_i > 0$) or not ($h_i < 0$). $\boldsymbol{g} = (g_{mj})_{m = 1,2, \ldots, M; j = 1,2, \ldots, S}$ is the effect of explicit environmental factors. $g_{mj}$ represents the effect of the $m$ th explicitly observed environmental factor on the $j$ th members. $\boldsymbol{J} = (J_{ij})_{i = 1,2, \ldots, S; j = 1,2, \ldots, S}$ is the species-species association matrix, where $J_{ij}$ represents the likelihood of co-occurrence between $i$ th and $j$ th species (large $J_{ij}$ indicates the coexistence of the two species more likely to occur). The energy attributed to each community composition, $E(\boldsymbol{\sigma}^{(k)}, \boldsymbol{\varepsilon})$, is computed based on those parameters using EPMEM. The occurrence probability of a community state $\boldsymbol{\sigma}^{(k)}$ is negatively proportional to $E(\boldsymbol{\sigma}(k), \boldsymbol{\varepsilon})$, and thus lower $E(\boldsymbol{\sigma}(k), \boldsymbol{\varepsilon})$ is more frequently occurred. In the landscape network, the energy gap between two proximal community compositions indicates directionality of state transitions (Fig. 1), whereby a lower energy state is more likely to occur than a higher energy state.

**SML performance affects the predictability in competitive Lotka-Volterra system**

Improving the parameter-fitting accuracy in EPMEM imply the better performance in overall ELA, yet it is still uncertain whether these approaches show improvements in the prediction of key properties in the landscape and global transitions of communities. To assess the performance of the fine-tuning method in ELA for capturing the emergent population dynamics by interspecies interactions, we employed the Lotka–Volterra (LV) model framework (Fig. S3) and tested whether the hyperparameter selection can increase the predictability in the population-level outcomes. The species-interaction parameter matrix $\boldsymbol{A}$ is randomly generated, and dynamics of 48 taxa were simulated in 2048 patches with random invasions until the distribution of the community compositions was converged (see Supporting Methods). The community compositions observed at the end point were regarded as stable states of each LV system. We focused on the LV system converged to more than 3 stable states and randomly sampled 512 points in the time-course data, which further converted to the species occurrence matrix (i.e., binary matrix) and used for the training dataset for the EPMEM parameter fitting.

Here, our aim is to check the correlation between the intrinsic stability of each LV system and the predicted landscape by ELA, and then the two-performance metrics, (1) stable states and (2) appearance probability of the community composition, were used as performance metrics for the overall ELA analysis (Fig. S3B). The predictability of the "stable states" was evaluated from the two aspects, "recall" (i.e., what fraction of the true stable states were recovered by ELA) and "precision" (i.e., what fraction of the predicted stable states corresponds to the true outcomes). Same as the case of parameter-fitting performance in EPMEM, recall values showed significant increase by the implementation of AdamW and Sparse modeling (Fig. S3C), indicating that those two algorithms enable ELA to recover the possible stable states more widely. In contrast, the precision value is significantly lower when including the AdamW and Sparse modeling, and thus those two methods harbored more false positives. This is primarily due to more sensitive estimation of the landscape, which facilitated the prediction of stable states that were not frequent in LV simulations (Fig. S3D). The second parameter, the "appearance probability", measures how often each community composition appears in the LV simulations. We then calculated the correlation between the empirical probabilities from the LV simulations, $\hat{P}(\sigma^{(k)})$ , and those predicted by EPMEM, $P(\sigma^{(k)})$. The implementation of AdamW and Sparse modeling also increased the correlation between those two parameters (Fig. S3E), indicating that those methods can facilitate the predictions for the stability more comprehensively.

### Supporting Methods

#### Formatting datasets

ELAplus requires the taxon abundance table (samples - taxa) and optionally, environmental parameters in the corresponding samples. In the `Formatting()` function, taxon abundances within each sample are first normalized to relative abundances and are converted into a binary presence/absence matrix by classifying taxa with relative abundances above a user-defined threshold as present (1) and all others as absent (0). Then, taxa with mean occurrence frequencies that are too low or too high are filtered out,  because rare taxa occur too infrequently to estimate reliable parameters, whereas ubiquitous taxa show little variation across samples and therefore contribute little to the inference of the energy landscape. If environmental factor data (e.g., temperature and pH) is available, such metadata are aligned with the samples of taxon abundance data and formatted for subsequent analyses.

#### Parameter fitting to the simulated datasets in EPMEM

Fitting the parameters of the EPMEM is performed using a stochastic approximation algorithm. We approximate the intractable model expectation in the likelihood gradient with a persistent (warm-started) Gibbs/heat-bath chain advanced by one sweep per parameter iteration, corresponding to persistent contrastive divergence with one sweep (PCD-1), also known as stochastic maximum likelihood (SML) as previously described [2]. SML estimates of the model parameters are obtained by minimizing the discrepancy between the sufficient statistics of the observed data and those implied by the model.

Let $\mathbf{Y} = (y_{ij}) \in \{0,1\}^{N \times S}$ represent a binary community state matrix, where each row represents a community state (sample) and each column represents a species (or taxa). The element $x_{ij} \in \{0,1\}$ indicates the absence or presence of species $j$ in sample $i$, and $\mathbf{E} = (\varepsilon_{ik}) \in \mathbb{R}^{N \times M}$ represent the environmental data matrix, where rows correspond to samples and columns correspond to environmental variables. The element $\varepsilon_{ik}$ denotes the value of environmental variable $k$ measured in sample $i$. SML repeatedly simulates data under the current parameters and then updates ***h**, **g**,* and ***J*** to reduce the discrepancies between observed vs simulated metrics. Two observed co-occurrence statistics are used for the fitting, $\mathbf{SS} = (\mathbf{Y})^{\mathrm{t}}\mathbf{Y}$ (here, $\mathbf{Y}^{\mathrm{t}}$ is the transpose of **Y**), and $\mathbf{SE} = (\mathbf{E})^{\mathrm{t}}\mathbf{Y}$. **SS** and **SE** are species–species co-occurrence and environment–species co-occurrence matrices, respectively. Then, we obtain the difference of sufficient statistics as:

$$\Delta\mathbf{SS} = \mathbf{SS} - \mathbf{SS}^* \tag{1}$$

and

$$\Delta\mathbf{SE} = \mathbf{SE} - \mathbf{SE}^* \tag{2}$$

$\mathbf{SS}^*$, $\mathbf{SE}^*$ are the statistics based on the $\mathbf{S}^*$, $\mathbf{E}^*$, which are simulated by EPMEM using the updated ***h**, **g**,* and ***J***.

Then, the algorith adjusts the model parameters to climb the approximate gradient, using a schedule of step sizes as:

$$\boldsymbol{h}^{(t)} = \boldsymbol{h}^{(t-1)} + \Delta\boldsymbol{h}^{(t)} \tag{3}$$

$$\boldsymbol{J}^{(t)} = \boldsymbol{J}^{(t-1)} + \Delta\boldsymbol{J}^{(t)} \tag{4}$$

$$\boldsymbol{g}^{(t)} = \boldsymbol{g}^{(t-1)} + \Delta\boldsymbol{g}^{(t)} \tag{5}$$

Here,

$$\Delta\boldsymbol{h}^{(t)} = \eta(1-m)\boldsymbol{G}_h + m\Delta\boldsymbol{h}^{(t-1)} \tag{6}$$

$$\Delta\boldsymbol{J}^{(t)} = \eta(1-m)\boldsymbol{G}_J + m\Delta\boldsymbol{J}^{(t-1)} \tag{7}$$

$$\Delta \boldsymbol{g}^{(t)} = \eta(1-m)\boldsymbol{G}_g + m\Delta \boldsymbol{g}^{(t-1)} \tag{8}$$

and,

$$\boldsymbol{G}_h = \frac{\mathrm{diag}(\Delta \mathbf{SS})}{N} \tag{9}$$

$$\boldsymbol{G}_J = \frac{\Delta \mathbf{SS}}{N} |\mathbf{I}(S) - 1| \tag{10}$$

$$\boldsymbol{G}_g = \frac{\Delta \mathbf{SE}}{N} \tag{11}$$

$\eta$ is the learning rate. $\boldsymbol{G}_h, \boldsymbol{G}_J, \boldsymbol{G}_g$ are the approximated likelihood gradients. Here, $\mathbf{I}(S)$ is a $S \times S$ identity matrix.

In the case where convergence was checked by the "update statistic", moving average of the **ΔSS** or **ΔSE** are used as metrics. Specifically, the mean of the most recent 100 values of the update metric was monitored, and the iteration was terminated when this moving average failed to improve (i.e., decrease) for more than 1000 consecutive iterations. Update statistic at *t* iteration and the convergence algorithm is defined as follows:

$$u_t = \frac{1}{2}\left| \frac{1}{NS^2}\sum_{i,j}(\Delta \mathbf{SS})_{ij} + \frac{1}{MNS}\sum_{i,j}(\Delta \mathbf{SE})_{ij} \right| \tag{12}$$

$$m_t = \frac{1}{L_t}\sum_{k=t-L_t+1}^{t} u_k \tag{13}$$

$$L_t = \min(t+1, 100) \tag{14}$$

$$\text{if } m_t < \min_{s<t} m_s \text{, then reset counter to 0; otherwise increment counter } ic \text{ by 1}$$

$$\text{if } ic > 1000 \rightarrow \text{stop}$$

**Implementation of AdamW algorithm**

We employ an adaptive moment estimation with weight decay (AdamW) optimizer to update the parameters [10]. The gradients are normalized and used to update first and second moment estimates, followed by bias correction and weight updates with decoupled weight decay. ***h**, **g**,* and ***J*** are updated as follows:

$$\boldsymbol{h}^{(t)} = \mathrm{diag}(\boldsymbol{\theta}_{\mathrm{SS}}) \tag{15}$$

$$\boldsymbol{J}^{(t)} = \boldsymbol{\theta}_{\mathrm{SS}} \text{, where } \mathrm{diag}(\boldsymbol{\theta}_{\mathrm{SS}}) = 0 \tag{16}$$

$$\boldsymbol{g}^{(t)} = \boldsymbol{\theta}_{\mathrm{SE}} \tag{17}$$

$\boldsymbol{\theta}_{\mathrm{SS}} = (\theta_{i,k}) \in \mathbb{R}^{S\times S}$ represent the species (taxa)-species (taxa) interaction matrix, where rows and columns correspond to species (or taxa). $\boldsymbol{\theta}_{\mathrm{SE}} = (\theta_{i,k}) \in \mathbb{R}^{S\times M}$ represent the species-environment interaction matrix, where rows correspond to species (or taxa) and columns correspond to environmental variables. Both matrices are updated by the same algorthm and denoted as $\boldsymbol{\theta}$ in the following explanation. $\boldsymbol{\theta}$ is updated in each step *t* as follows.

$$\boldsymbol{\theta}^{(t)} = \boldsymbol{\theta}^{(t-1)} + \eta \frac{\widehat{\mathbf{m}}^{(t)}}{\sqrt{\hat{v}_t} + \varepsilon} - \eta w \boldsymbol{\theta}^{(t-1)} \tag{18}$$

where,

$$\mathbf{m}^{(t)} = (1-\beta_1)\Delta\boldsymbol{\theta}^{(t)} + \beta_1 \mathbf{m}^{(t-1)} \tag{19}$$

$$\widehat{\mathbf{m}}^{(t)} = \frac{\mathbf{m}^{(t)}}{1-\beta_1^{(t)}} \tag{20}$$

$$\mathbf{v}^{(t)} = (1-\beta_2)\left(\Delta\boldsymbol{\theta}^{(t)}\right)^2 + \beta_2 \mathbf{v}^{(t-1)} \tag{21}$$

$$\hat{\mathbf{v}}^{(t)} = \frac{\mathbf{v}^{(t)}}{1-\beta_2^{(t)}} \tag{22}$$

$$\Delta\boldsymbol{\theta}_{\mathrm{SS}}{}^{(t)} = \frac{\Delta\mathbf{SS}^{(t)}}{N} \tag{23}$$

$$\Delta\boldsymbol{\theta}_{\mathrm{ES}}{}^{(t)} = \frac{\Delta\mathbf{SE}^{(t)}}{N} \tag{24}$$

The updated parameter $\beta_1^{(t)}$ and $\beta_2^{(t)}$ are also used for AdamW, which updatd at each iteration $t$ as

$$\beta_1^{(t)} = \beta_1\, \beta_1^{(t-1)} \tag{25}$$

$$\beta_2^{(t)} = \beta_2\, \beta_2^{(t-1)} \tag{26}$$

and *w* is the weight decay in adamW algorithm.

To prevent the model from overfitting the training data and encourage sparsity in species interactions $\boldsymbol{J}$, we apply an L1-type regularization after each gradient step based on that applied to stochastic gradient descent [19]. This procedure shrinks small interaction coefficients toward zero while preserving the diagonal terms, $\boldsymbol{h}$. The L1-type regularization is applied to the parameters after the update by momentum or adamW, and the parameters updated by those methods at step $t$ is designated as

$$\boldsymbol{J}^{\left(t-\frac{1}{2}\right)} = \left(J_{ij}^{\left(t-\frac{1}{2}\right)}\right) \tag{27}$$

$$\text{If } J_{ij}^{\left(t-\frac{1}{2}\right)} > 0, \qquad J_{ij}^{(t)} = \max\left(0, J_{ij}^{\left(t-\frac{1}{2}\right)} - \left(u^{(t)} + q_{ij}^{(t)}\right)\right) \tag{28}$$

$$\text{If } J_{ij}^{\left(t-\frac{1}{2}\right)} < 0, \qquad J_{ij}^{(t)} = \min\left(0, J_{ij}^{\left(t-\frac{1}{2}\right)} + \left(u^{(t)} - q_{ij}^{(t)}\right)\right) \tag{29}$$

$$\text{If } J_{ij}^{\left(t-\frac{1}{2}\right)} = 0 \text{ or } i = j, \qquad J_{ij}^{(t)} = 0 \tag{30}$$

where,

$$q_{ij}^{(t)} = \sum_{k}^{t-1}\left(J_{ij}^{(k)} - J_{ij}^{\left(k-\frac{1}{2}\right)}\right) \tag{31}$$

$$u^{(t)} = \eta\lambda N t \text{ (adamW)} \tag{32}$$

$$u^{(t)} = 0.1\eta\lambda N t \text{ (momentum)} \tag{33}$$

$\lambda$ is the strength of the penalty.

**Evaluation of model predictive performance**

To evaluate predictive performance, we calculated the mean negative conditional log-likelihood of observed species occurrences under the fitted model. For each sample $i$ and species $s$, we computed the conditional probability of occurrence as

$$\hat{p}_{is} = \frac{1}{1 + \exp[-(h_s + \sum_{m=1}^{M} \varepsilon_{im} g_{ms} + \sum_{t=1}^{S} y_{it} J_{ts})]} \tag{34}$$

The occurrence probability assigned by the model to the observed state ($\mathbf{Y} = (y_{ij}) \in \{0,1\}^{N\times S}$) was then defined as

$$q_{is} = |1 - y_{is} - \hat{p}_{is}| \tag{35}$$

and predictive performance was quantified by

$$\mathcal{L}_{\text{val}} = -\frac{1}{NS}\sum_{i=1}^{N}\sum_{s=1}^{S}\log q_{is} \tag{36}$$

Lower values of $\mathcal{L}_{\text{val}}$ indicate better predictive performance. In `Findbp()`, the function performs the grid-search of hyper-parameters: $\eta$: learning rate, $\lambda$: the strength of L1 penalty, $w$: the weight decay parameter, and the total iterations of algorithm.

**Generation of random EPMEM parameters and occurrence datasets**

To evaluate parameter-estimation accuracy, we generated binary occurrence datasets from randomly specified EPMEM parameters. Each intrinsic occurrence parameter in $\boldsymbol{h}$ was independently drawn from a uniform distribution ranging from −2 to 2, denoted as $\boldsymbol{U}(-2, 2)$, hereafter. The target connectance of the pairwise parameter matrix ($\boldsymbol{J}$), defined as the proportion of nonzero taxon pairs, was set to either 0.2 or 0.5. Taxon pairs were randomly selected according to the specified connectance, and their parameters were independently drawn from $\boldsymbol{U}(-2, 2)$. Here, 2 environmental variables were included, and each taxon-specific environmental coefficient in $\boldsymbol{g}$ was independently drawn from $\boldsymbol{U}(-2, 2)$. These generating values of $\boldsymbol{h}$, $\boldsymbol{J}$, and $\boldsymbol{g}$ were treated as the true parameters when evaluating estimation accuracy.

For each parameter set, an occurrence dataset containing 512 samples was generated using heat-bath Gibbs sampling. All samples were initialized as all-absent states. The environmental variables for each sample ($\boldsymbol{\varepsilon}$) were independently drawn from $\boldsymbol{U}(0, 1)$ and held constant throughout sampling.

At each heat-bath update, one taxon was selected at random for each sample and its state was resampled as present or absent according to the probability $p_{is}$, while the states of all other taxa remained unchanged. For each sample $i$ and taxon $s$, $p_{is}$ was calculated as:

$$p_{is} = \frac{1}{1 + \exp[-(h_s + \sum_{m=1}^{M}\varepsilon_{im}g_{ms} + \sum_{t=1}^{S}x_{it}J_{ts})]} \tag{37}$$

Here, $x_{it} \in \{0,1\}$ denotes the current occurrence state of taxon $t$ in sample $i$.

This procedure was repeated for 500 iterations, and the final states of the 512 sampling chains were used as the simulated occurrence matrix.

The EPMEM was then fitted to each simulated dataset, and parameter-estimation accuracy was quantified using Spearman correlations between the generating (true) and estimated values.

**Simulating population dynamics of competitive Lotka-Volterra model**

To test the applicability of our method to community assembly dynamics driven by population-level processes, we used the following Lotka–Volterra (LV) equation:

$$\frac{dx_i}{dt} = x_i \left( r_i - \sum_j a_{ij} x_j \right) \tag{38}$$

Here, $x_i$ represents the population abundance of species (taxa) $i$ in the abundance vector $\mathbf{x}$, and $r_i$ denotes the intrinsic growth rate of species $i$, while $a_{ij}$ represents the interaction coefficient between species $i$ and $j$ (i.e., interspecies interactions), an element of an interaction matrix $\boldsymbol{A} = (a_{ij})$. Here, we tested the case of competitive Lotka-Volterra system, and $a_{ij}$ is always negative. To simulate community dynamics, we introduced the process of "extinction" and "recruitment" into our model. Extinction is defined as the frequency of species $i$ decreasing below a certain threshold (i.e., population size $x_i < 10^{-5}$). Recruitment is defined as the introduction of new species into the system at $r = 10^{-4}$.

We generated a LV dataset that characterizes the stability landscape of a LV competition model. First, we generated 2048 sites and randomly sampled the interaction strengths ($\boldsymbol{A}$), initial population structure ($\mathbf{x}_0 = (x_i) \in \{0, r\}^S$ , $r$ is the propagule size), and recruitment processes independently. The number of species, $S$, was fixed throughout the simulation. We obtained the numerical solutions of the LV equation by integrating the differential equation until equilibrium was reached. We used the first-order Euler method with $dt$ = 0.2. The algorithm checked the species extinction, then randomly selected one of them, and introduced it with a propagule size at $r = 10^{-4}$ at every 500 steps. During these processes, the abundance vector $\mathbf{x}$ of the LV model was recorded every 100 invasions. We applied a binarization process to the abundance vectors. Specifically, species were treated as present (1) if $x_i > 0.001$, and absent (0) otherwise. The resulting presence–absence vectors $\boldsymbol{\sigma}^{(k)}$ were then used as the community composition. Let $\hat{p}(\boldsymbol{\sigma}^{(k)})$ be the present probability of community compositions of 2048 sites and $\hat{q}(\boldsymbol{\sigma}^{(k)})$ be the probability at previous recording point (i.e., 100 invasions ago). To check the convergence of the population dynamics, we calculated the Jensen-Shannon divergence between them as:

$$D_{JS} = \sqrt{\frac{\left\{\sum \hat{p}(\sigma^{(k)}) \log \frac{\hat{p}(\sigma^{(k)})}{\mu_{pq}(\sigma^{(k)})} + \sum \hat{q}(\sigma^{(k)}) \log \frac{\hat{q}(\sigma^{(k)})}{\mu_{pq}(\sigma^{(k)})}\right\}}{2}} \tag{39}$$

The simulation was stopped when the $D_{JS}$ fell below 0.0001.

**Pruning stable states**

For each pair of stable states $i$ and $j$, `ELPruning()` calculates their energy barriers relative to the tipping point connecting them as follows:

$$\Delta E_i = E_{tip} - E_i \tag{40}$$

$$\Delta E_j = E_{tip} - E_j \tag{41}$$

The pair is assigned a priority value:

$$\Delta E_{ij} = min(\Delta E_i, \Delta E_j) \tag{42}$$

This represents the depth of its shallower basin. For each state, a bootstrap-approximated p-value is calculated as the proportion of bootstrap replicates in which its barrier is negative. All pairs are ordered by $\Delta E_{ij}$ and the first pair for which either p-value exceeds the specified threshold is selected. Within the selected pair, the higher-energy state, which has the shallower basin, is merged into the lower-energy state. The stable-state and tipping-point matrices are then updated, and the procedure is repeated until no pair satisfies the pruning criterion.

**Constructing disconnectivity graph**

Disconnectivity graphs were generated from the inferred energy landscape to summarize the relationships among stable states. For each pair of stable states, the minimum energy barrier was calculated as the lowest tipping-point energy along the minimum-energy path connecting the two states. A hierarchical tree was then constructed by progressively joining stable states according to these barrier heights, such that the vertical axis represents energy and branch heights correspond to transition barriers between basins. This representation provides a compact visualization of the multi-stable structure of the energy landscape and facilitates comparison of basin depths and transition pathways.

### Supporting Figures

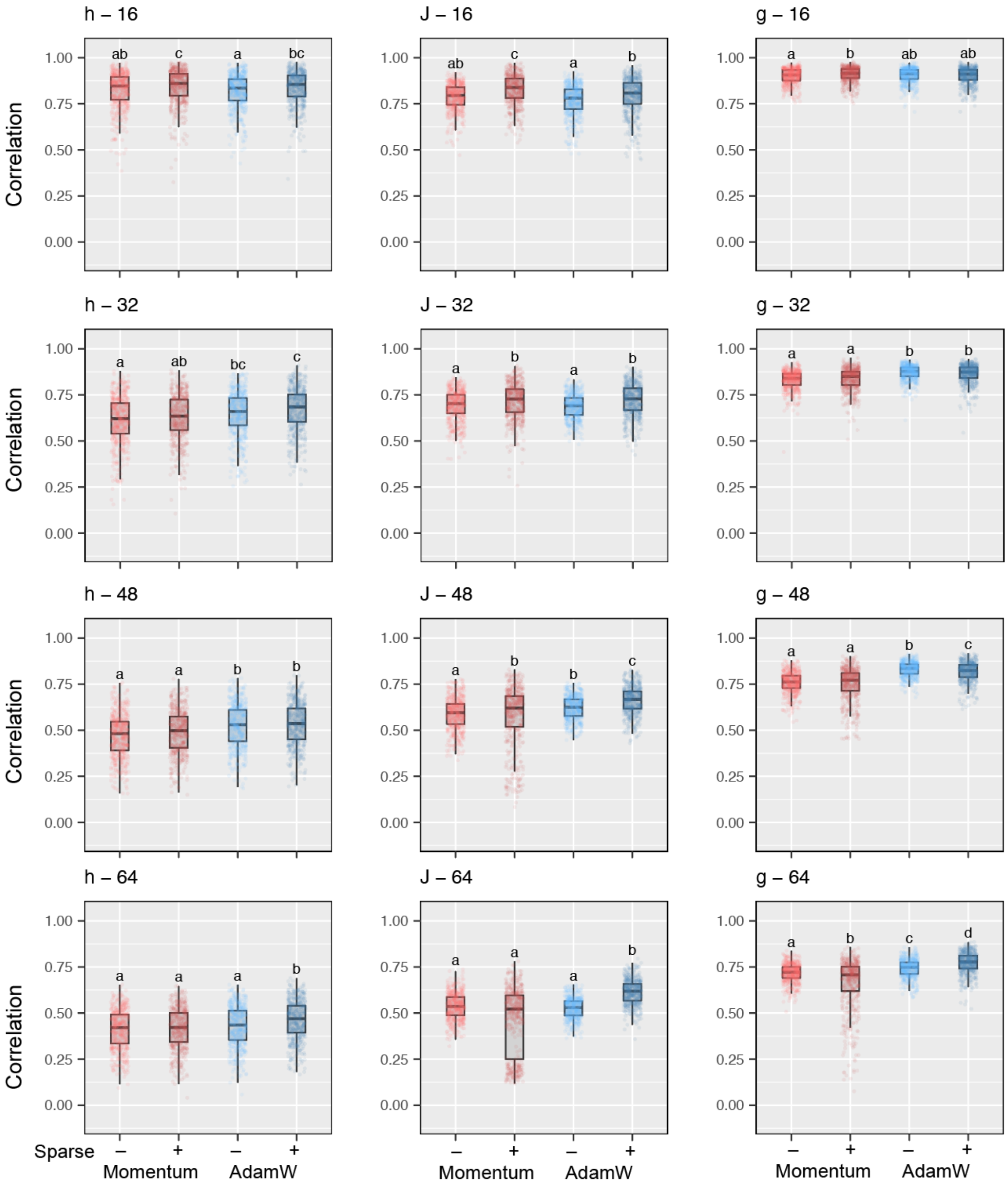


**FIGURE S1. Effects of AdamW and sparse modeling on low-connectance community composition data.** Spearman's correlations between true and predicted EPMEM parameters (***h***, ***J***, and ***g***) for 512 random parameter sets. Numbers above each panel indicate the number of taxa in the randomly generated parameter sets. Bold lines indicate median values, and statistically significant groups are denoted by different letters. The connectance of taxa–taxa interactions (i.e., the proportion of nonzero off-diagonal elements in ***J***) was set to 0.2.

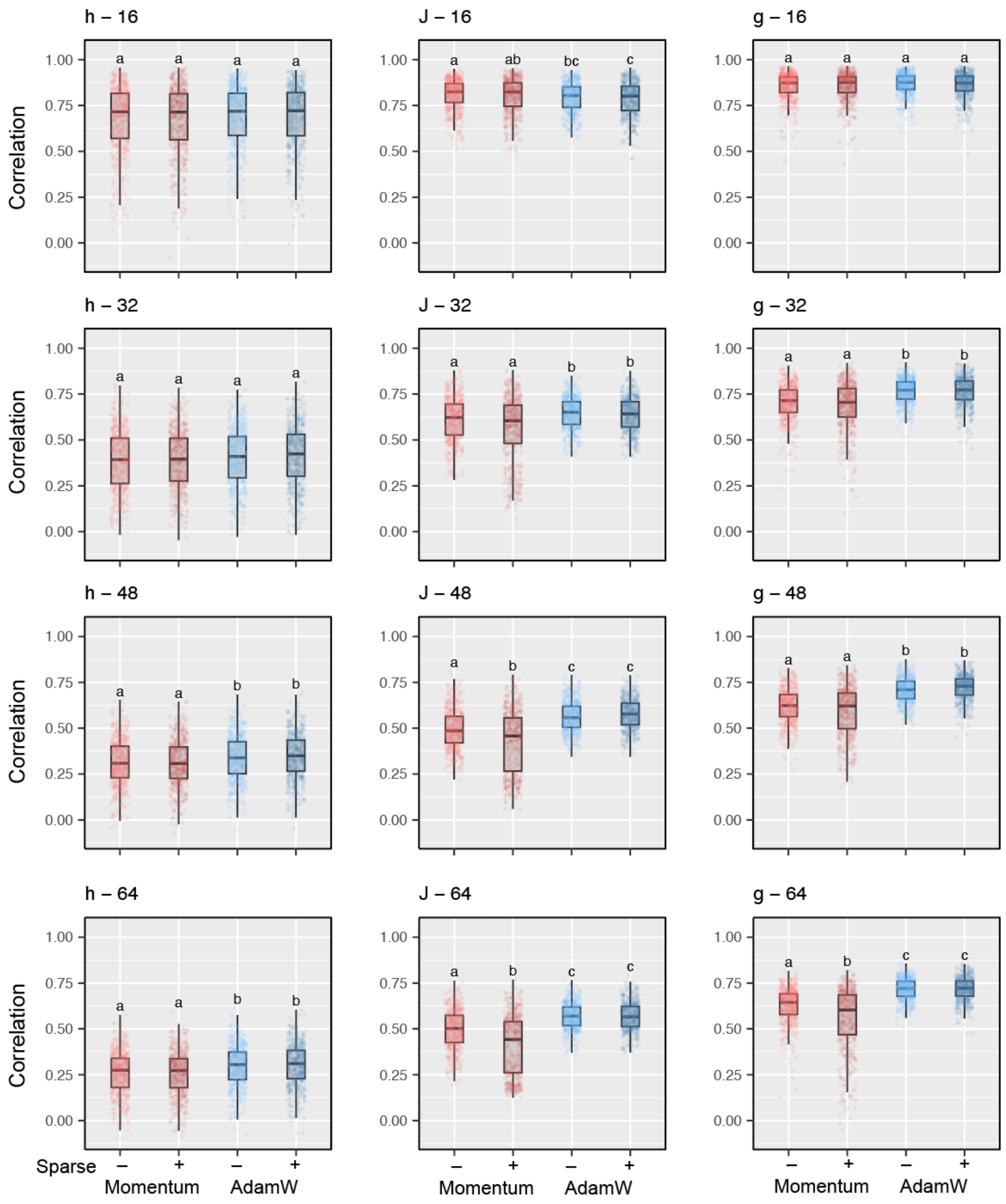


**FIGURE S2. Implementation of AdamW and sparse modeling improved predictability in the fitting performance.** Spearman's correlations between true and predicted parameter sets in EPMEM (***h***, ***J***, and ***g***) for 512 random parameter sets. Simulations, parameter fitting, and validation were performed under the same conditions as in Fig. S1 and Fig. 2, except that the connectance of taxa–taxa interactions (i.e., the proportion of nonzero off-diagonal elements in ***J***) was set to 0.5. The number indicated above each panel shows the number of taxa in the randomly generated parameters. Bold lines indicate median values, and statistically significant groups are denoted by different letters**.**

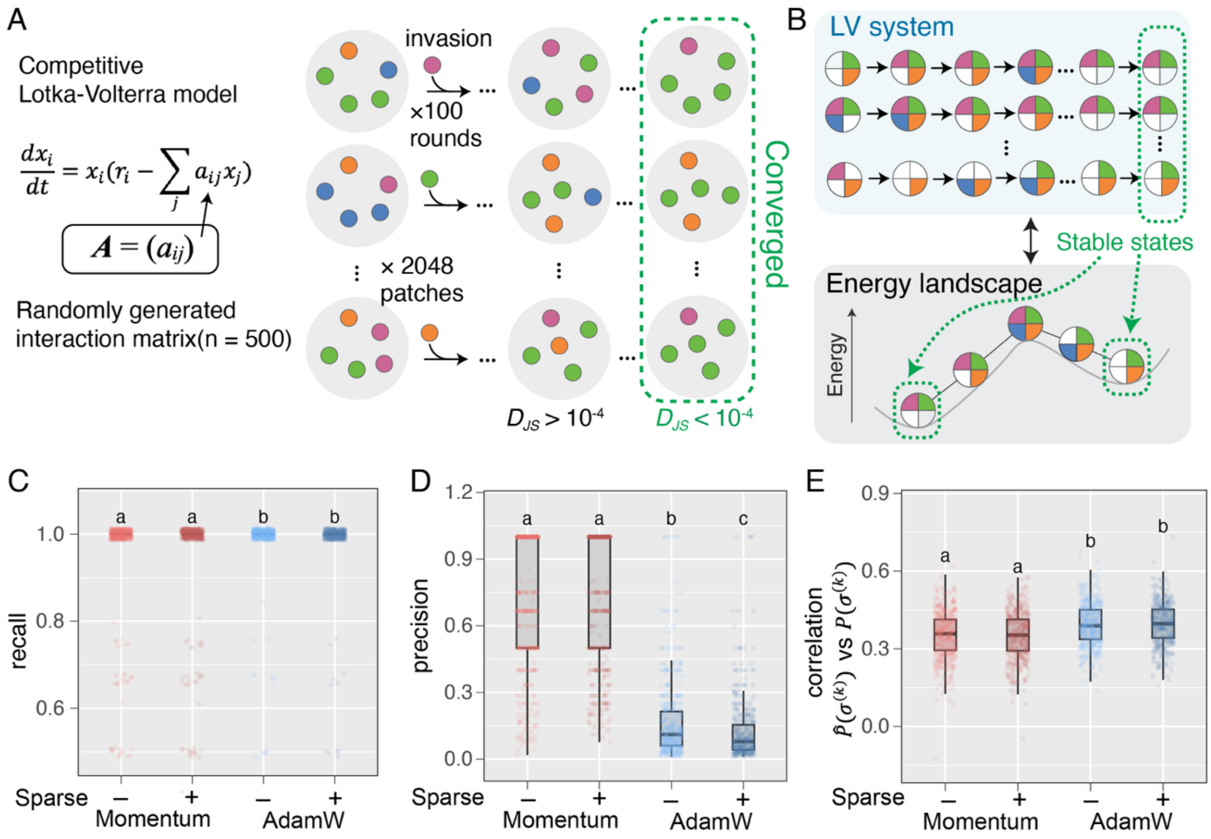


**FIGURE S3. Additional optimization algorithms improve prediction accuracy in the energy landscape analysis.** (A) Overview of the competitive Lotka–Volterra (LV) simulations. Species interaction matrices ***A*** were randomly generated for 500 parameter sets to comprehensively simulate population dynamics. For each matrix, 2048 patches with different initial community compositions were simulated. Random invasions were repeated until the Jensen–Shannon divergence ($D_{JS}$) of community composition distributions across patches converged ($D_{JS} < 10^{-4}$). (B) Schematic comparison between the LV system and ELA predictions. Final community compositions after convergence in the LV simulations were regarded as stable states. Compositions observed in more than 1% of patches (> 20 patches) were defined as “true” stable states and compared with those predicted by ELA. (C) Recall of stable states predicted by ELA relative to those observed in the LV simulations. (D) Precision of stable states predicted by ELA. (E) Spearman’s correlations between empirical appearance probabilities in the LV simulations and those predicted by ELA. In panels C–E, bold lines indicate median values, and statistically significant groups are denoted by different letters.
